# Medical Nonadherence, Cannabis Use, and Renal Outcome in Systemic Lupus Erythematosis

**DOI:** 10.1101/389973

**Authors:** Basmah A. Jalil, Clifford R. Qualls, Romy J. Cabacungan, Wilmer L. Sibbitt, James I. Gibb, Leonard E. Noronha, Roderick A. Fields, N. Suzanne Emil, Monthida Fangtham, Arthur D Bankhurst

## Abstract

**Background/Objective:** Non-adherence to recommended medical therapy has been associated with poorer outcomes in systemic lupus erythematosus (SLE). The present research investigated the association of medical non-adherence and cannabis use on renal outcomes of SLE.

**Methods:** This was a prospective 5-year longitudinal outcome study of 276 female SLE patients 30.4% who chronically used medical cannabis and 69.5% who did not. Outcomes were determined at 5 years after enrollment in the study.

**Results:** Cannabis use in SLE patients was associated with an increased prevalence of neuropsychiatric SLE (p<0.05), opioid analgesic use (p<0.01), cigarette smoking (p<0.001), and non-adherence to the medical regimen (non-cannabis: 3% non-adherence vs. cannabis use: 95% non-adherence, p<0.001). Within the 5-year period, the cannabis group demonstrated a 53% increase in mortality (p=0.12) and 127% increase in end-stage renal disease requiring dialysis (p<0.001). With logistic regression analysis adjusting for SLE disease activity (SLEDAI-2K), cannabis use was an independent predictor of end-stage renal disease: Odds ratio 2.65 (CI 1.32 – 5.32, p<0.01). Adjusting for SLE disease damage (SLICC/ACR-DI), cannabis use remained an independent predictor of end-stage renal disease: Odds ratio 2.0 (CI 1.26 – 6.23, p<0.01). With multivariable analysis adjusting for non-adherence, the effect of cannabis on end-stage renal disease could be largely attributed to an increase in non-adherence to medical therapy.

**Conclusions:** Non-adherence to recommended therapy and medical cannabis use are associated with a significant increase in the development of end-stage renal disease in SLE.

## Background/Purpose

Medical cannabis has been used to treat recalcitrant cancer pain, anorexia, and the nausea of chemotherapy [1]. Cannabis has also been used to treat non-cancer pain, including the pain of sensory neuropathy, multiple sclerosis, fibromyalgia, mixed chronic pain, and rheumatoid arthritis [1-3]. Certain medical practitioners incorporate medical cannabis into the therapy of osteoarthritis, rheumatoid arthritis and systemic lupus erythematosus (SLE) with the goal of preventing opioid use, avoiding non-steroidal anti-inflammatory drug toxicity, preserving renal function, and reducing corticosteroid use [3,4]. Based on extensive review of the relevant medical literature, Allan and colleagues have recently published guidelines for medical cannabis that recommend that medical cannabis be restricted to resistant medical conditions for which there is some evidence of beneficial effect (neuropathic pain, palliative and end-of-life pain, chemotherapy-induced nausea and vomiting, and spasticity due to multiple sclerosis or spinal cord injury) (1,2). Although there has been important research into the effects of medical cannabis in many other conditions characterized by chronic pain, the literature for cannabis-based management in SLE is especially lacking [1-4].

In the present study, we examined the 5-year outcomes of SLE patients who regularly used medical cannabis versus those who did not use cannabis. The present research, to our knowledge, is one of the first epidemiologic investigations into associations of medical cannabis use with SLE disease outcomes, including end-stage renal disease.

## Methods

This research was approved by the institutional review board (IRB) and was in adherence with the Helsinki Declaration and subsequent revisions. The study design was a cross-sectional and longitudinal observational study of outcome. 276 SLE female patients were studied over a period of 5 years and were enrolled consecutively as encountered in clinic. Each subject provided written informed consent. Inclusion criteria included any patient with SLE, age 18-80. Exclusion criteria were age < 18 years, age > 80 years, male gender, pre-existing end-stage renal disease, renal transplantation, and any patient with an autoimmune diagnosis other than SLE (overlap disease). The population sample included diverse social, racial, and ethnic backgrounds with North American Hispanics (58%) and Caucasian Whites (34%) being the dominant participants.

The diagnosis of SLE was established in each subject using the American College of Rheumatology (ACR) revised criteria for the classification for SLE [5]. SLE disease activity was determined with the System Lupus Erythematosus Disease Activity Index 2000 (SLEDAI-2K) and SLE disease injury was measured with the Systemic Lupus Erythematosus Collaborating Clinics/American College of Rheumatology Damage Index (SLICC/ACR-DI) [6,7]. Each of these metrics was further subcategorized into Neuro-SLEDAI-2K consisting of the neurologic components of SLEDAI-2K, and Neuro-SLICC/ACR-DI consisting of the neurologic components of SLICC/ACR-DI. Neuropsychiatric SLE (NPSLE) was characterized by the ACR nomenclature and case definitions for NPSLE [8]. After collection this prospective 5-year database was then de-identified. Additional IRB approval was obtained prior to analyzing the de-identified database in relation to medical cannabis use. Outcomes were determined at 5 years after enrollment in the study.

Medical cannabis in this study was defined as smoked cannabis (marijuana) and was categorically defined as at least 2 smoked cannabis units (cigarettes or pipes) per week and the subject was categorized as “cannabis” or “non-cannabis” with 84 patients (30.4%) regularly using cannabis and 192 (69.5%) with no cannabis use. New Mexico is one of the states in the USA with locally legal medical cannabis programs. Medical cannabis was not prescribed or provided by the rheumatologist, rather by the patient’s primary care provider, medical cannabis specialist, or pain specialist. Demographics and outcomes recorded were age of SLE onset (years), age (years), SLE disease duration (years), tobacco use (years), tobacco use (pack per day), tobacco use ever, opiate use, weight (kilograms), pain, morning stiffness (hours), SLICC/ACRDI, Neuro-SLICC/ACRDI, Non-Neuro-SLICC/ACRDI, SLEDAI-2K, Neuro-SLEDAI-2K, Non-Neuro-SLEDAI-2K, antinuclear antibody (ANA) titer, anti-DNA antibody (IU), anti-phospholipid antibodies (IgG, IgM, IgA in GPL, MPL, APL units, respectively), rheumatoid factor (IU), antiribosmal P (IU), family history (arthritis, SLE, rheumatoid arthritis), active renal disease (glomerulonephritis), renal failure/end-stage renal disease, night pain, death, and Sjogren’s syndrome. Pain was accessed with the 10 cm visual analogue pain scale (VAS). Opioid dosage (the study population predominately used oxycodone 5-10 mg tablets) was normalized to 7.5 mg morphine milligram equivalent (MME) (equivalent to 5 mg oxycodone), and was reported as the number of 7.5 mg MME tablets per month [9]. Non-adherence (non-adherence) was defined by admission by the patient that they were not taking their SLE medications, blood testing for mycophenolate levels, and/or by missing more than 20% of their clinic appointments (“a no-show”); cancellations with rescheduling and hospitalizations were not counted as “no-show” or non-adherence. Survival and death of SLE subjects were determined by following up with each patient or, if the patient could not be contacted, the patient’s family, autopsy reports from the medical examiner, chart review, and searching the Social Security

Death Index for death/benefit files relating to the study participant [10]. This combined methodology permitted 100% follow-up of all subjects at 5 years. Renal failure/end-stage renal disease was defined in a patient who 1) required long-term hemo-or peritoneal dialysis, or 2) required and underwent renal transplantation.

### Statistical Methods

The major comparison was between cannabis and non-cannabis users. Statistical differences between measurement data were determined with Student t-test and categorical data with Fisher’s exact method, associations were determined initially with univariable regression analysis and afterward multivariable models were created to determine independent effects on dependent variables.

## Results

Demographics and results of the 276 subjects with SLE are shown in Tables 1 and 2. No significant differences were observed in SLE disease duration, SLE disease activity (SLEDAI-2K), injury/damage from SLE (SLICC/ACR-DI), ANA titer, dsDNA antibody, anti-phospholipid antibodies (IgG, IgM, IgA), rheumatoid factor, active renal disease, night pain, morning stiffness, pain by VAS, Sjogren’s syndrome, anti-Smith antibody, RNP antibody, and anti-ribosomal P antibody in the cannabis and non-cannabis SLE groups.

**TABLE 1.**
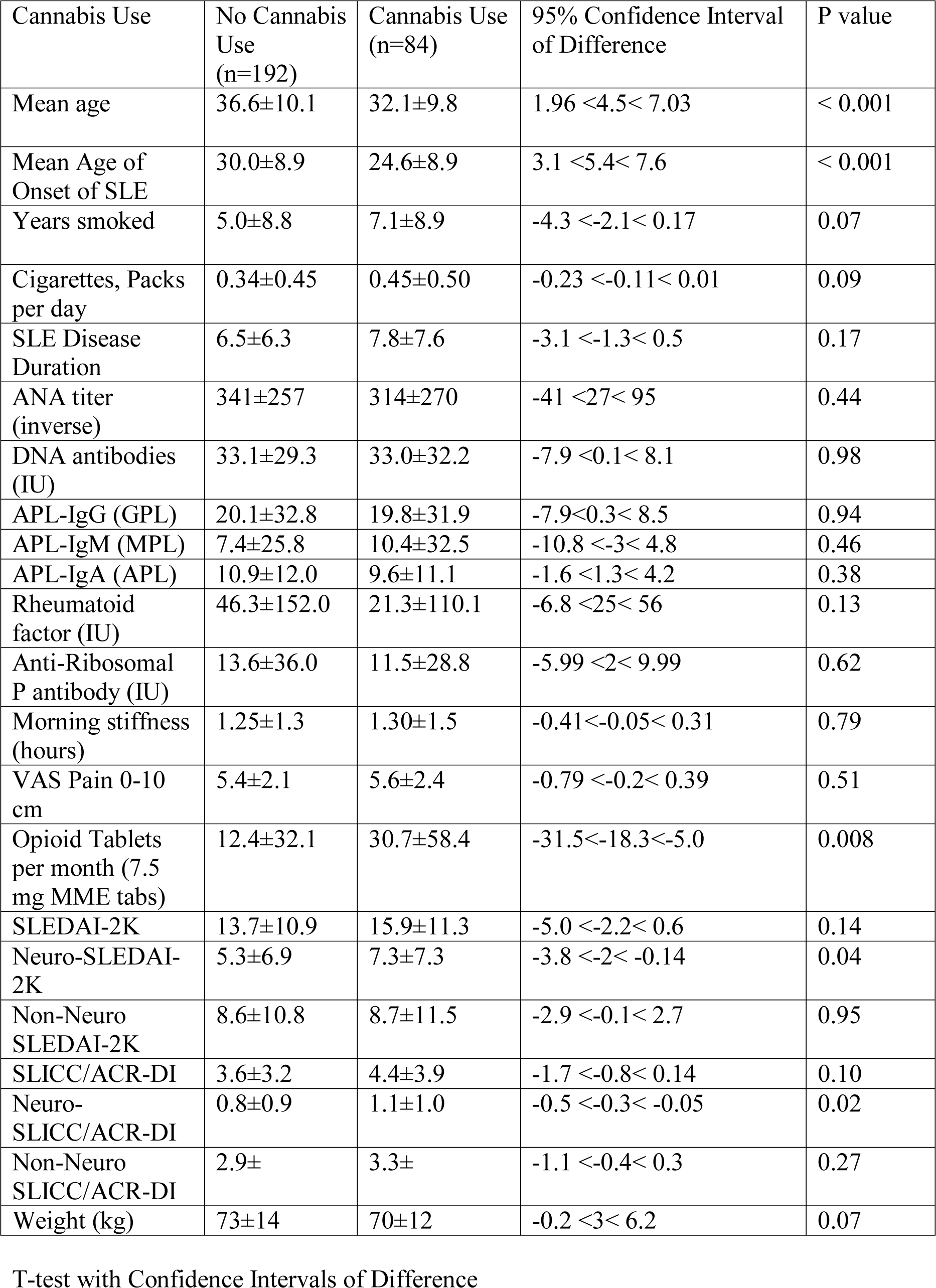
Comparison of SLE Patients with no Cannabis Use versus Cannabis Use.

| Cannabis Use | No Cannabis Use<br>(n=192) | Cannabis Use<br>(n=84) | 95% Confidence Interval<br>of Difference | P value |
| --- | --- | --- | --- | --- |
| Mean age | 36.6±10.1 | 32.1±9.8 | 1.96 <4.5< 7.03 | < 0.001 |
| Mean Age of<br>Onset of SLE | 30.0±8.9 | 24.6±8.9 | 3.1 <5.4< 7.6 | < 0.001 |
| Years smoked | 5.0±8.8 | 7.1±8.9 | -4.3 <-2.1< 0.17 | 0.07 |
| Cigarettes, Packs<br>per day | 0.34±0.45 | 0.45±0.50 | -0.23 <-0.11< 0.01 | 0.09 |
| SLE Disease<br>Duration | 6.5±6.3 | 7.8±7.6 | -3.1 <-1.3< 0.5 | 0.17 |
| ANA titer<br>(inverse) | 341±257 | 314±270 | -41 <27< 95 | 0.44 |
| DNA antibodies<br>(IU) | 33.1±29.3 | 33.0±32.2 | -7.9 <0.1< 8.1 | 0.98 |
| APL-IgG (GPL) | 20.1±32.8 | 19.8±31.9 | -7.9<0.3< 8.5 | 0.94 |
| APL-IgM (MPL) | 7.4±25.8 | 10.4±32.5 | -10.8 <-3< 4.8 | 0.46 |
| APL-IgA (APL) | 10.9±12.0 | 9.6±11.1 | -1.6 <1.3< 4.2 | 0.38 |
| Rheumatoid<br>factor (IU) | 46.3±152.0 | 21.3±110.1 | -6.8 <25< 56 | 0.13 |
| Anti-Ribosomal<br>P antibody (IU) | 13.6±36.0 | 11.5±28.8 | -5.99 <2< 9.99 | 0.62 |
| Morning stiffness<br>(hours) | 1.25±1.3 | 1.30±1.5 | -0.41<-0.05< 0.31 | 0.79 |
| VAS Pain 0-10<br>cm | 5.4±2.1 | 5.6±2.4 | -0.79 <-0.2< 0.39 | 0.51 |
| Opioid Tablets<br>per month (7.5<br>mg MME tabs) | 12.4±32.1 | 30.7±58.4 | -31.5<-18.3<-5.0 | 0.008 |
| SLEDAI-2K | 13.7±10.9 | 15.9±11.3 | -5.0 <-2.2< 0.6 | 0.14 |
| Neuro-SLEDAI-2K | 5.3±6.9 | 7.3±7.3 | -3.8 <-2< -0.14 | 0.04 |
| Non-Neuro<br>SLEDAI-2K | 8.6±10.8 | 8.7±11.5 | -2.9 <-0.1< 2.7 | 0.95 |
| SLICC/ACR-DI | 3.6±3.2 | 4.4±3.9 | -1.7 <-0.8< 0.14 | 0.10 |
| Neuro-SLICC/ACR-DI | 0.8±0.9 | 1.1±1.0 | -0.5 <-0.3< -0.05 | 0.02 |
| Non-Neuro<br>SLICC/ACR-DI | 2.9± | 3.3± | -1.1 <-0.4< 0.3 | 0.27 |
| Weight (kg) | 73±14 | 70±12 | -0.2 <3< 6.2 | 0.07 |

T-test with Confidence Intervals of Difference

**TABLE 2.** Comparison of SLE Patients with and without Cannabis Use

| Cannabis Use | No Cannabis Use<br>(n=192) | Cannabis Use<br>(n=84) | P value |
| --- | --- | --- | --- |
| Sjogren's Syndrome | 51.0% | 51.2% | 0.98 |
| Family history SLE | 30.2% | 44.1% | 0.03 |
| Tobacco use ever<br>(cigarettes) | 34.9% | 58.5% | < 0.001 |
| Active Alcohol Use | 5.2% | 10.7% | 0.12 |
| Night pain | 52.4% | 55.9% | 0.60 |
| Joint pain | 78.0% | 69.0% | 0.13 |
| Active<br>glomerulonephritis | 19.8% | 20.2% | 1.00 |
| Non-adherence | 2.1% | 92.8% | <0.001 |
| Death within 5 years | 15.6% | 23.8% | 0.12 |
| End-stage Renal<br>Disease within 5<br>years | 11.0% | 25.00% | 0.006 |
| Opioid Analgesic<br>Use | 19.8% | 34.5% | 0.03 |

However, the cannabis smokers had a younger age (p<0.001), a younger age of SLE onset (p<0.001), a more common family history of SLE (p=0.03), a greater percentage of opioid analgesic use (p=0.01), more consumption per month of 7.5 mg MME tablets of opioid analgesics (p=0.008), increased tobacco use (p<0.001), increased active neuropsychiatric SLE symptoms (Neuro-SLEDAI-2K) (p=0.04), more neurologic injury (Neuro-SLICC/ACR-DI) (p=0.02), increased non-adherence with medical therapy (non-cannabis non-adherence: 3% vs. cannabis use: 95% non-adherence, p value < 0.001), and over a 5 year period a 127% increase in end-stage renal disease (non-cannabis: 11% vs. cannabis: 25%, p=0.006).

The percentage of deaths over a 5 year period increased in the cannabis group by 53% (non-cannabis: 15.6% deaths/5 years; cannabis: 23.8% deaths/5 years,) although this did not reach statistical significance (p=0.12)

Logistic regression analysis adjusting for disease activity (SLEDAI-2K) cannabis use was an independent predictor of end-stage renal disease: Odds ratio 2.65 (CI 1.32 – 5.32, p= 0.006).

Adjusting for disease injury (SLICC/ACR-DI) cannabis use remained an independent predictor of end-stage renal disease: Odds ratio 2.0 (CI 1.26 – 6.23, p= 0.01).

With multivariable analysis adjusting for non-adherence cannabis use was not an independent predictor of end-stage renal disease: Odds ratio 0.9 (CI 0.7 – 1.1, p= 0.92). Similarly, with multivariable analysis adjusting for cannabis use non-adherence was also not an independent predictor of end-stage renal disease: Odds ratio 1.0 (CI 0.8 – 1.2, p= 0.94). Thus, with multivariable analysis adjusting for both cannabis use and non-adherence, the effect of cannabis use on the increase in end-stage renal disease could be statistically explained by an increase in non-adherence to the recommended medical regimen. Thus, cannabis use and non-adherence were statistically dependent on each other and were essentially equal predictors of developing end-stage renal disease over a 5 year period.

## Discussion

The present study demonstrates that medical cannabis use in SLE is not associated with lower VAS pain scores, lower SLE disease activity or injury, preserved renal function, or other beneficial findings, but rather is associated with increased neuropsychiatric SLE, opioid analgesic use, cigarette smoking, and non-adherence with the recommended medical regimen with a 53% increase in the death rate and a 127% increase in end-stage renal disease within the 5 year period (Tables 1 and 2). SLE is often characterized by chronic pain related to arthritis, neuropathy, Raynaud’s phenomenon, low back pain, vertebral compression fractures, headache, fibromyalgia, and aseptic necrosis of the hips [11]. Although many SLE patients can cope with their pain, a substantial proportion (up to 42%) do not cope well and show signs of pain-related dysfunction, including distress, activity interference, catastrophising, and interpersonal difficulties [12]. Treating pain in these individuals is difficult due to concomitant coagulation and renal disorders often excluding the use of non-steroidal anti-inflammatory drugs, thus, there has been a recent interest in treating chronic pain in SLE with medical cannabis to avoid the potential toxic effects of non-steroidal anti-inflammatory drugs, opiates, and excessive corticosteroids [4].

Cannabinoids are a group of compounds present in the Cannabis plant (Cannabis sativa L.) and mediate their physiological and behavioral effects by activating specific cannabinoid receptors [1-3]. With the discovery of the cannabinoid receptors (CB1 and CB2) and the endocannabinoid system, research in this field has expanded exponentially. Cannabinoids have been shown to act as potent immunosuppressive and anti-inflammatory agents in vitro and have been reported to mediate potential beneficial effects in a wide range of immune-mediated diseases such as multiple sclerosis, diabetes, peripheral neuropathy, septic shock, rheumatoid arthritis, and allergic asthma [1-3]. Thus, since cannabinoids can be immunosuppressive and there are cannabinoid receptors in symptomatic tissues, it might be expected that cannabis use might reduce the activity of SLE and improve SLE outcomes.

The prototypical use of smoked medical cannabis is in cancer therapy to reduce anxiety, reduce nausea, improve appetite, and reduce pain, where cannabis is not expected to substantially alter the outcome of the disease [1,2]. In other chronic diseases with less increased mortality than cancer, such as chronic low back pain, osteoarthritis, myofascial-fibromyalgia, and impingement neuropathy, the positive or negative effects of cannabis outcome would require extremely large cohorts with many years of observation to determine significant differences in mortality and morbidity since these diseases themselves have only long-term effects on mortality [1,2,13]. However, without effective medical therapy SLE has a much accelerated morbidity and mortality than many of these other chronic diseases, thus the beneficial or deleterious effects of medical cannabis on outcome if present could potentially be observed in a much shorter time frame [14].

Similar to the results of the present study in SLE patients, the use of cannabis in other diseases has been associated with an increased rate of non-adherence to medications and recommended medical therapy [15-17]. In human immunodeficiency virus (HIV) patients cannabis use has been associated with a nearly 100% increase in non-adherence to antiretroviral therapy [15]. Cannabis use has also been associated with non-adherence to insulin regimens in diabetes mellitus resulting in increased hospitalizations for ketoacidosis and infections [16]. Cannabis use is also associated with non-adherence to medications in depression and other psychiatric conditions [17]. In the present study in SLE patients, medical cannabis use was associated with an increase to 93% non-adherence to the recommended medical regimen (Table 2), further supporting the close association of cannabis and non-adherence reported in other conditions [15-17]. This increase in non-adherence to the medical regimen with cannabis use in SLE is of even greater concern since the baseline non-adherence to medical therapy in SLE is reported also to be very high, ranging from 43% to 83% [18,19].

Using Medicaid data Feldman and colleagues have reported that SLE patients are up to 79-83% non-adherent with medication and that this non-adherence is associated with greater use of emergency services and increased hospitalizations [20]. Uribe, Segovia, and colleagues have demonstrated that non-adherence in SLE was associated with a younger age, single marital status, ‘no shows’ to clinics, greater disease activity and more severe SLE manifestations (serositis, renal involvement, positive anti-ds-DNA antibodies) that are associated with poorer outcomes [21]. Rivera et al reported that non-adherence to medications in SLE patients is associated with increased flares of lupus, including active lupus glomerulonephritis [22]. Renal transplantation often utilizes similar immunosuppressive drug regimens that are used to treat lupus nephritis, and non-adherence to immunosuppressive therapy has been increasingly recognized as a major contributing cause to episodes of rejection and graft loss [23,24]. Thus, since cannabis use has been associated with non-adherence to therapy, and non-adherence to therapy has been associated with more severe SLE manifestations and active lupus glomerulonephritis, it is not surprising that cannabis use is also associated with increased non-adherence and an associated increased prevalence of end-stage renal disease in SLE as shown in the present study (Table 2) [15-24].

It is possible that cannabis use contributes to non-adherence to medical regimens in SLE by the well-known deleterious effects of cannabinoids on motivation, memory, and other cognitive functions [13-17,25]. Cannabis use has previously been associated with increased neurocognitive defects in autoimmune diseases, a finding that the present study confirms with increased prevalence of neuropsychiatric SLE (increased Neuro-SLEDAI-2K and increased Neuro-SLICC/ACR-DI) (Table 1) [25]. Further, cannabis use in musculoskeletal diseases and arthritis has been associated with the increased use of narcotic analgesics, which the present study confirms in SLE (Tables 1 and 2) [26]. The present study demonstrated no beneficial changes in immune parameters, SLE disease activity (SLEDAI-2K), or SLE disease damage (SLICC/ACRDI) in medical cannabis users, but increases in neuropsychiatric SLE (Neuro-SLEDAI-2K and Neuro-SLICC/ACRDI) and increased end-stage renal disease suggesting that the immunomodulatory effects of medical cannabis in SLE are either minor or are overwhelmed by the marked increase in non-adherence to recommended medical therapy (Tables 1 and 2).

Patients with SLE have a high risk for glomerulonephritis that without effective medical intervention often results in renal failure and the need for dialysis and renal transplantation [14,27,28]. Cannabis use has been reported to cause membranous glomerulopathy, another reason for caution in SLE patients with preexisting renal disease [29]. In contrast, Greenan and colleagues have reported that recreational cannabis use was not associated with worse outcomes after renal transplantation; however, most of these patients were not SLE patients [30]. To date there are few published trials regarding renal transplant survival specifically in SLE patients who do and do not utilize medical cannabis, and the present study does not provide further information on this group.

The present study demonstrates that end-stage renal disease increases by 127% in SLE patients who are medical cannabis users, and that the statistical cannabis effect can be explained by an increase in non-adherence with the medical regimen. Medical cannabis use is also associated with increased neuropsychiatric symptoms in SLE patients, and it is also possible that deterioration in cognition and motivation exacerbated by cannabis may be contributing to increased non-adherence to therapy, a factor that may be important in that a majority of SLE patient already have baseline neurocognitive defects [8,25]. Presently, guidelines for the use of medical cannabis in SLE have not been formulated largely because there is little objective data on which to base these recommendations. However, based on the present study and a review of the published medical literature, SLE is presently not one of the conditions where medical cannabis can be administered with expectations of a predictable benefit [1,2].

There are a number of limitations to this study. The association of cannabis and non-adherence with poorer outcomes may reflect greater disease activity, severity, and individual predisposition rather than be directly causative. Medical cannabis use could predispose to the observed non-adherence to therapy, but an alternative is that the addictive personality type who uses medical cannabis is the same personality type who is non-adherent to therapy with or without cannabis. This latter possibility is supported by the increase in concomitant substance use (tobacco and opioid analgesics) in the medical cannabis users (Tables 1 and 2). Secondly, this study was restricted only to women with SLE, not men, as men with SLE generally have a poorer outcome and different drug use patterns than women with SLE. However, most SLE patients are women, so this study is generally applicable to the majority of SLE patients. The present study was a cross-sectional, prospective cohort study that accurately reflected our SLE population over 5 years, but was not an incipient cohort study with subjects enrolled on diagnosis and was not a randomized controlled trial, thus, the reported associations may not be causative. The period of observation was 5 years, longer or shorter periods of observation may have different results. Since the potency and regimens for use of medical cannabis have not been standardized, the dosing and the potency of the medical cannabis were intrinsically variable, thus, certain SLE patients could have had extremely potent or weak cannabis that could have influenced the results. Finally, only smoked medical cannabis was studied, not capsules, tablets, topical preparations, extracts, synthetic cannabinoids, or purified cannabinoids that are increasingly available.

## Conclusion

This 5-year outcome study indicates that medical cannabis use in SLE is not associated with reduced pain, less use of opioid analgesics or diminished SLE disease activity. Rather, medical cannabis use in SLE is associated with an increased incidence of neuropsychiatric SLE, accentuated opioid use, increased non-adherence to recommended therapy, and an increased progression to end-stage renal disease.

## REFERENCES

1. Allan GM, Ramji J, Perry D, Ton J, Beahm NP, Crisp N, Dockrill B, Dubin RE, Findlay T, Kirkwood J, Fleming M, Makus K, Zhu X, Korownyk C, Kolber MR, McCormack J, Nickel S, Noël G, Lindblad AJ. Simplified guideline for prescribing medical cannabinoids in primary care. Can Fam Physician. 2018 Feb;64(2):111–120.

2. Allan GM, Finley CR, Ton J, Perry D, Ramji J, Crawford K, Lindblad AJ, Korownyk C, Kolber MR. Systematic review of systematic reviews for medical cannabinoids: Pain, nausea and vomiting, spasticity, and harms. Can Fam Physician. 2018 Feb;64(2):e78–e94.

3. Wright S, Ware M, Guy G. The use of a cannabis-based medicine (Sativex) in the treatment of pain caused by rheumatoid arthritis. Rheumatology (Oxford). 2006 Jun;45(6):781; author reply 781-2. Epub 2006 Apr 18. PMID:16621924

4. Líndal E, Thorlacius S, Stefánsson JG, Steinsson K. Pain and pain problems among subjects with systemic lupus erythematosus in Iceland. Scand J Rheumatol. 1993;22(1):10–3. PMID: 8434240

5. Hochberg MC. Updating the American College of Rheumatology revised criteria for the classification of systemic lupus erythematosus. Arthritis Rheum 1997 Sep;40(9):1725. PMID:9324032

6. Gladman DD, Ibañez D, Urowitz MB. Systemic lupus erythematosus disease activity index 2000. J Rheumatol. 2002 Feb;29(2):288–91.

7. Gladman D, Ginzler E, Goldsmith C, Fortin P, Liang M, Urowitz M, et al. The development and initial validation of the Systemic Lupus International Collaborating Clinics/American College of Rheumatology damage index for systemic lupus erythematosus. Arthritis Rheum. 1996 Mar;39(3):363–9. PMID:8607884

8. American College of Rheumatology. The American College of Rheumatology nomenclature and case definitions for neuropsychiatric lupus syndromes. Arthritis Rheum. 1999 Apr;42(4):599–608. PMID:10211873

9. Centers for Disease Control and Preventiona. CDC Guideline for Prescribing Opioids for Chronic Pain. https://www.cdc.gov/drugoverdose/prescribing/guideline.html accessed 09/29/2017.

10. U.S., Social Security Death Index, 1935–2014. Ancestry.com. Original data: Social Security Administration. Social Security Death Index, Master File. Social Security Administration, USA. http://search.ancestry.com/search/db.aspx?dbid=3693, accessed Sept.25, 2017.

11. Di Franco M, Guzzo MP, Spinelli FR, Atzeni F, Sarzi-Puttini P, Conti F, Iannuccelli C. Pain and systemic lupus erythematosus. Reumatismo 2014 Jun 6;66(1):33–8. doi: 10.4081/reumatismo.2014.762. PMID:24938194

12. Fischin J, Chehab G, Richter JG, Fischer-Betz R, Winkler-Rohlfing B, Willers R, Schneider M. Factors associated with pain coping and catastrophising in patients with systemic lupus erythematosus: a cross-sectional study of the LuLa-cohort. Lupus Sci Med. 2015 Nov 12;2(1):e000113. doi: 10.1136/lupus-2015-000113. eCollection 2015 PMID:26629351

13. Davstad I, Allebeck P, Leifman A, Stenbacka M, Romelsjö A. Self-reported drug use and mortality among a nationwide sample of Swedish conscripts - A 35-year follow-up. Drug Alcohol Depend. 2011 Nov 1;118(2-3):383–90. doi: 10.1016/j.drugalcdep.2011.04.025. Epub 2011 Jun 12. PMID:21664771

14. Tektonidou MG, Lewandowski LB, Hu J, Dasgupta A, Ward MM. Survival in adults and children with systemic lupus erythematosus: a systematic review and Bayesian meta-analysis of studies from 1950 to 2016. Ann Rheum Dis. 2017 Aug 9. pii: annrheumdis-2017-211663. doi: 10.1136/annrheumdis-2017-211663. [Epub ahead of print] PMID: 28794077

15. Gross IM, Hosek S, Richards MH, Fernandez MI. Predictors and Profiles of Antiretroviral Therapy Adherence Among African American Adolescents and Young Adult Males Living with HIV. AIDS Patient Care STDS. 2016 Jul;30(7):324–38.

16. Isidro ML, Jorge S. Recreational drug abuse in patients hospitalized for diabetic ketosis or diabetic ketoacidosis. Acta Diabetol. 2013 Apr;50(2):183–7. doi: 10.1007/s00592-010-0243-z. Epub 2010 Dec 7. PMID:21136122

17. Stewart SL, Baiden P. An exploratory study of the factors associated with medication nonadherence among youth in adult mental health facilities in Ontario, Canada. Psychiatry Res. 2013 May 30;207(3):212–7. doi: 10.1016/j.psychres.2013.01.017. Epub 2013 Mar 5. PMID:23465295

18. Mehat P, Atiquzzaman M, Esdaile JM, AviÑa-Zubieta A, De Vera MA. Medication Nonadherence in Systemic Lupus Erythematosus: A Systematic Review. Arthritis Care Res (Hoboken). 2017 Nov;69(11):1706–1713.

19. Feldman CH, Collins J, Zhang Z, Subramanian SV, Solomon DH, Kawachi I, Costenbader KH. Dynamic patterns and predictors of hydroxychloroquine nonadherence among Medicaid beneficiaries with systemic lupus erythematosus. Semin Arthritis Rheum. 2018 Jan 8. pii: S0049-0172(17)30376-1. doi: 10.1016/j.semarthrit.2018.01.002 [Epub ahead of print]

20. Feldman CH, Yazdany J, Guan H, Solomon DH, Costenbader KH. Medication Nonadherence Is Associated With Increased Subsequent Acute Care Utilization Among Medicaid Beneficiaries With Systemic Lupus Erythematosus. Arthritis Care Res (Hoboken). 2015 Dec;67(12):1712–21

21. Uribe AG, Ho KT, Agee B, McGwin G Jr, Fessler BJ, Bastian HM, Reveille JD, Alarcón GS. Relationship between adherence to study and clinic visits in systemic lupus erythematosuspatients: data from the LUMINA cohort. Lupus. 2004;13(8):561–8.

22. Rivera F, Anaya S. Lupus nephritis flare in young patients: relapse or nonadherence to treatment? Int J Nephrol Renovasc Dis. 2014 Mar 27;7:117–21.

23. Nevins TE, Nickerson PW, Dew MA. Understanding Medication Nonadherence after Kidney Transplant. J Am Soc Nephrol. 2017 Aug;28(8):2290–2301.

24. Chand S, Atkinson D, Collins C, Briggs D, Ball S, Sharif A, Skordilis K, Vydianath B, Neil D, Borrows R. The Spectrum of Renal Allograft Failure. PLoS One. 2016 Sep 20;11(9):e0162278. doi: 10.1371/journal.pone.0162278. eCollection 2016.

25. Markusse HM, van den Bent MJ, Vecht CJ. Immunology in medical practice. XIV. Central nervous system complications in systemic autoimmune diseases Ned Tijdschr Geneeskd. 1998 Mar 7;142(10):508–12. PMID:9623096

26. Fleming MF, Balousek SL, Klessig CL, Mundt MP, Brown DD. Substance use disorders in a primary care sample receiving daily opioid therapy. J Pain. 2007 Jul;8(7):573–82. Epub 2007 May 11. PMID:17499555

27. Croca SC, Rodrigues T, Isenberg DA. Assessment of a lupus nephritis cohort over a 30-year period. Rheumatology (Oxford). 2011 Aug;50(8):1424–30. doi: 10.1093/rheumatology/ker101. Epub 2011 Mar 16. PMID: 21415024

28. Franco C, Yoo W, Franco D, Xu Z. Predictors of end stage renal disease in African Americans with lupus nephritis. Bull NYU Hosp Jt Dis. 2010;68(4):251–6. PMID:21162701

29. Bohatyrewicz M, Urasinska E, Rozanski J, Ciechanowski K. Membranous glomerulonephritis may be associated with heavy marijuana abuse. Transplant Proc. 2007 Dec;39(10):3054–6. PMID:18089320

30. Greenan G, Ahmad SB, Anders MG, Leeser A, Bromberg JS, Niederhaus SV. Recreational marijuana use is not associated with worse outcomes after renal transplantation. Clin Transplant. 2016 Oct;30(10):1340–1346.

